# A Bayesian approach to multivariate and multilevel modelling with non-random missingness for hierarchical clinical proteomics data

**DOI:** 10.1101/153049

**Authors:** Irene SL Zeng, Thomas Lumley, Katya Ruggierol, Martin Middleditch

## Abstract

High throughput mass-spectrometry-based proteomics data from clinical studies brings challenges to statistical analysis. The challenges originate from the hierarchical levels of protein abundance data and interactions between clinical study design and experimental design. The non-random missingness of the measurements from a vast amount of information also adds complexity in data analysis. We propose multivariate multilevel models to analyse protein abundances and to handle abundance-dependent missingness within a Bayesian framework. The proposed model enables the variance decomposition at different levels of the data hierarchy and provides shrinkage of protein-level estimates for a group of proteins. A logistic missingness and censored model with informative prior is used to handle incomplete data. Hamiltonian MC/No-U-Turn Sampling and Gibb MCMC algorithms are created to derive the posterior distribution of study parameters; Hamiltonian MC is demonstrated to gain more efficiency for these high-dimensional correlated data. Improvements of the proposed missing data model is compared to the univariate mixed effect model and the multivariate-multilevel model using complete data in a simulated study and a clinical proteomics study. The proposed model framework can be used in other types of data with similar structure and Non Random Missingness mechanism (MNAR).

## 1. Introduction

### 1.1. Clinical Proteomic studies

Proteomics has been used as a system biology technique in pharmaceutical industry and medical research to investigate hundreds and thousands of molecular biomarkers simultaneously. It belongs to the family of system biology, including metabolomics, transcriptomics and genomics. These “omic” fields study interacting molecular networks inter-disciplinarily. Like aforementioned “omics”, proteomics utilizes biotechnology platforms that can systematically identify proteins and quantify their abundances. One of the popular platforms used for protein discovery is mass spectrometry, which can accurately determine the molecular mass of ions for peptide/protein sequencing and quantitation [1-4]. Through different ionization and detection processes, mass spectrometry (MS) coupled with other techniques (i.e. chromatography) can separate different peptide species from a biological sample and produce a large amount of ion intensity data [5-9]. These intensity data are used to inform the abundances of the peptides and from these to construct the abundances of the corresponding proteins. The ***hierarchical structure*** of the protein abundance originates from the biochemical process of the mass spectrometry-based proteomic experiment [10]. When a proteomic experiment involves more than one biological sample, through assay batching or multiplexing [11-14], multiple biological samples can be analysed simultaneously in a single-run experiment. A multiplex comprises a batch of multiple samples. Each sample is labelled in the multiplex for its identity in a mass spectrometry experiment. When multiple runs are involved, the biological samples will be allocated into *labels* and *runs* according to an experimental design. The statistical analysis of mass spectrometry (MS) data curated from multiple-run experiments of multiplex-assays requires us to understand the data structure fully. Firstly, due to biochemical processes (trypsin digestion and protein fragmentation), the abundance data is provided at the peptides level, not proteins. The proteins abundances must be derived from their observed constituent peptides’ abundances. Secondly, each protein is not identified by the same set of peptides in each experiment. Thirdly, the MS experimental factors could have different impacts across different peptides/proteins. Fourthly, a clinical proteomics study has two phases of study design: clinical study design and biological experimental design. The confounding effects from a MS-based experiment hence are multileveled. Using a multilevel model will be an advanced solution to coping these confounding variables.

### 1.2. Data structure and sources of variation of a Mass Spectrometry-based isotopic labelling experiment

The protein expression data curated from MS-based proteomics experiments can either be the relative or absolute abundance of proteins. The relative abundances of proteins are measured by the intensity of ions from the mass spectrometry, while the absolute abundance of the protein is measured by the concentration [10]. The relative quantitation of protein abundance utilize different labelling techniques including chemical, biological, metabolic and enzymatic incorporation in a multi-sample proteomics experiment [10-12]. One common labelling experiment is to use iTRAQ, an isobaric chemical labelling approach that is widely adopted in proteomics laboratories. iTRAQ is a biochemical reagent comprising a reactive group, a tag (label) with a mass balance group and a reporter group [10-12, 14-18]. The reporter groups of the iTRAQ reagent, which have different molecular mass-to-charge ratios ranging from 114.1-117.1 Th in the 4-plex assay and 113.1-121.1 Th in the 8-plex assay, are used to distinguish multiple samples in the assay. The intensities of the labelled peptides are used to inform the relative abundance of proteins [19].

### 1.3. The instrumental feature of the mass spectrometer and the quantification

In addition to the hierarchical structure of protein abundance data, the physical features of mass spectrometry also influence the variations in ions intensities. We discovered from the two proteomic iTRAQ studies that, the logarithmic intensity decreases with the mass-to-charge ratio and the molecular mass of the peptides. The identified variations in intensity measurements caused by MS’s physical natures in our case studies are consistent with the findings of the other studies [20, 21]. Breitwieser et al. (2011)[20] observed more variabilities for the lower intensity signals and illuminated a funnel-shaped association between the peptide ratio and the logarithmic intensities. This evidence suggested the relative abundance of a protein is likely to be underestimated by its large peptide components. The quantification of a protein thus needs to take into account the variations from instruments. The proposed multilevel model includes mass-to-charge ratios as a variable contributing to this variation and is shown to increase the precision of estimates for the unknown parameters.

### 1.4. The non-random missingness mechanism in data from mass spectromety

The missingness mechanisms for intensities data comprises both Missing-Not-At-Random (MNAR) and censoring. The MNAR mechanism of protein abundance is related to their peptides intensities, masses, electrical charges, and ionization efficiency[21, 22]. The likelihood of missing values thus may depend on the unknown protein abundance and the observable mass-to-charge ratio. The left-censored phenomenon is caused by the detecting limit of a MS. We model the probability of having a missing intensity as a function of the intensity and the mass-to-charge ratio [figure 1]. Data from mass spectrometry of smaller molecules (metabolomics) supports this model [21]. Their data showed that higher probabilities of having missing intensities were associated with observed lower abundance. There is a nonlinear relationship between probability of having noisy intensity value and the mass-to-charge ratio.

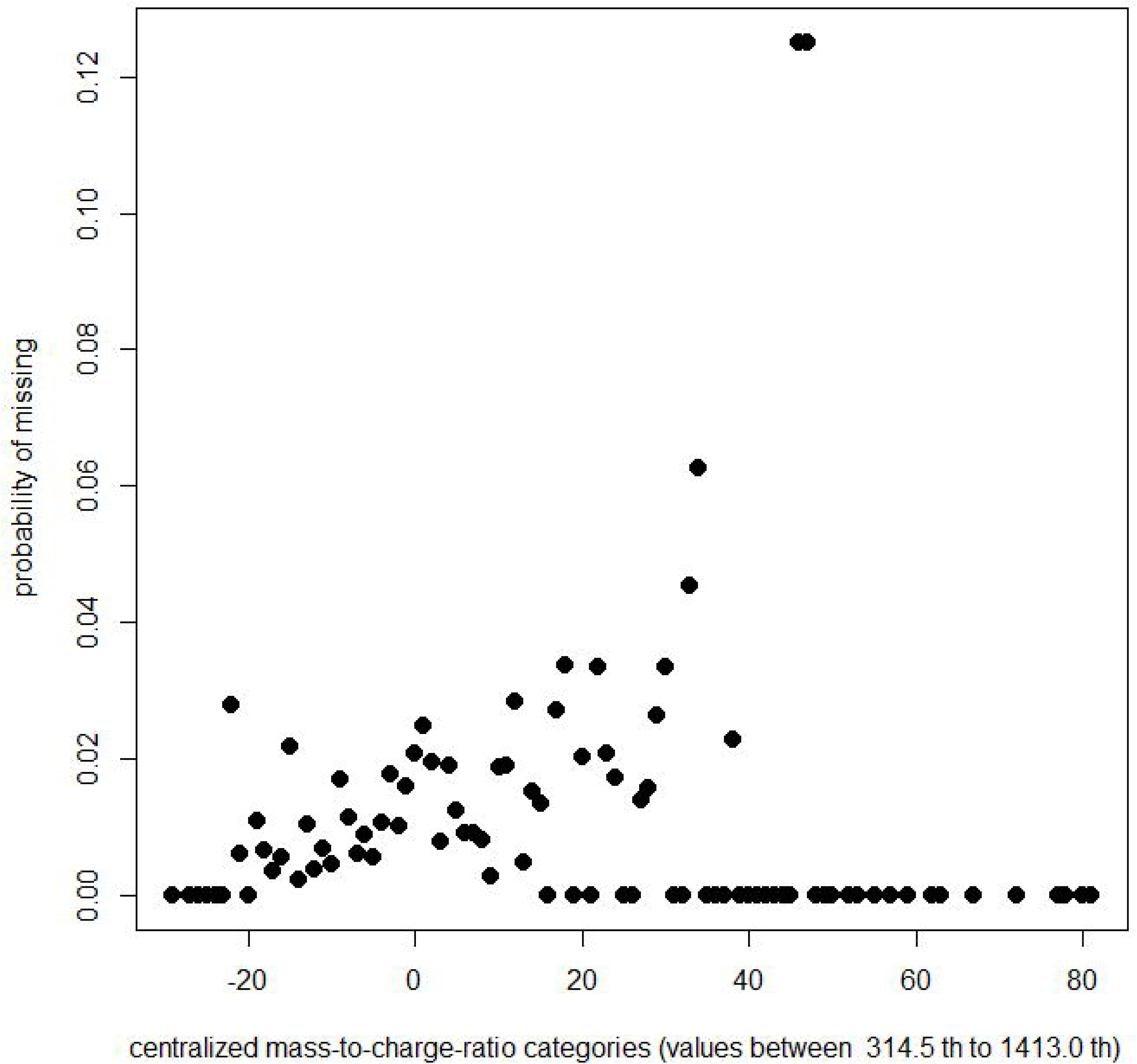

### 1.5. An imputation model under a Bayesian multivariate and multilevel inference framework

We use an inference model within a multivariate-multilevel framework to analyze the hierarchical protein abundance data generated by clinical proteomics studies. These models decompose variations from different sources, such as experimental factors, biological features of proteins and physiological characteristics of biological samples. We further model non-random missingness as either left-censored or completely missing. Little (1993), Little and Rubin (1987) [23, 24] proposed three likelihood-based methods (selection model, pattern-mixture model and shared-parameter model) for MNAR. Selection model, in which the missing data mechanism and parameters of the inferential model are conditionally estimated given the hypothetical complete data, are the appropriate approach for the abundance-dependent missing data. The hypothetical complete data comprises the missing and observed values. The known physical properties of peptides and mass spectrometers provide the **auxiliary information** for the missing data. We use the selection model factorization [24] for MNAR including the left-censored missingness under a Bayesian framework. We estimate the likelihood of missingness by using auxiliary variable mass-to-charge ratio (m/z) and the intensity values. The censored and non-censored missing intensities are marginalized for the complete data inferential likelihood, i.e.

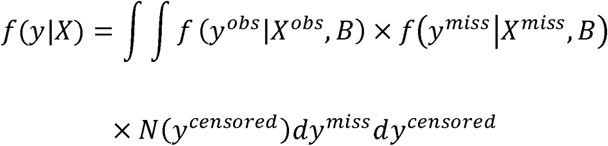
where *y* = (*y^obs^*, *y^miss^*, *y^censored^*), *y^obs^*, *y^miss^* and *y^censored^* represent the observed, missing and censored intensity values respectively; *X^obs^* and *X^miss^* are the corresponding explanatory variables used in the multivariate multilevel inferential model *f* for the observed and missing intensities. *N*(*y^censored^*) is the normal truncated function for the left-censored values. The parameters *B* of the multivariate multilevel inference model *f*, the missing values and the parameters of the missing mechanism are treated as unknowns, and are jointly estimated under a Bayesian framework. The Bayesian approach enables utilization of the prior information learned from the other studies, i.e. Hrydziuszko and Viant (2012) [21], as an enrichment of the missing value imputation.

## 2. A multivariate-multilevel model for a group of proteins without missing intensity

We begin by defining the multivariate framework, initially for the case of complete proteomic data.

In contrast to the traditional single-protein model popularly applied in analysis of proteomics datasets [25], the multiple proteins model analyzes a group of proteins within a multivariate and multilevel framework. Through the use of a two-level model, the multivariate model can be analyzed as a univariate model. In this two-level model, different proteins are treated as level 2 units, and peptide represented by reporter ions are considered the level one units nested within proteins and subjects in the data hierarchy. Ideally the same protein will be observed in all subjects; in such case, proteins and subjects are defined as crossed factors and both are the level 2 units. Although this is in general not the case in practice, the multilevel model allows for unbalances in the design and random missingness [26]. The level one part of the model defines the relationship between the peptides intensities and the experimental factors (MS experiment factors: i.e. Label and *n*th run). The level two part of the model describes the relationship between the selected protein level random coefficients and the explanatory variables. Another component of the level 2 model defines the relationship between a random intercept for subject and the subject level explanatory variables (clinical design factors: i.e. randomization group and baseline measures).

### Define level one part of the 2-level model

A full model with interactions at level one is defined as:

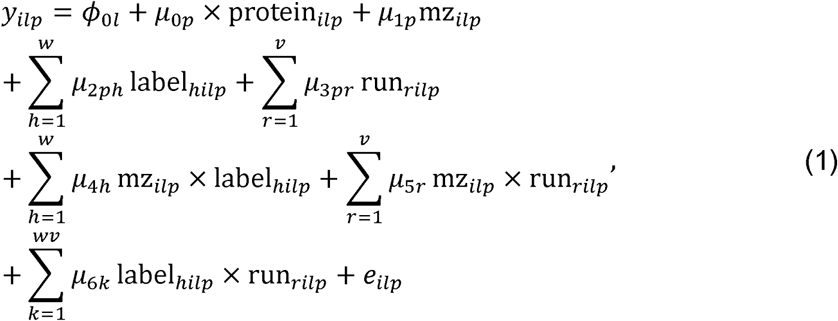

where *y_ilp_* denotes the log-transformed intensity for peptide *i* (*range* l: *n*;), subject *l* (*range* 1: *q*) and protein *p* (*range* 1: *z*). *protein_ilp_* denotes the protein identity for *p*; *mz_ilp_* denotes the m/z ratio for peptide *i*, subject *l* and protein *p*; *label_hilp_* is a binary variable with value 0 or 1 which indicates response *y_ilp_* is from label *h* (*range*1: *w*); *run_rilp_* is a binary variable with value 0 or 1 which indicates response *y_ilp_* is from run *r* (*range*1: *v*) of all experiments; *ϕ*_0*l*_ denotes the random intercept for subject *l*; *U =* (*μ*_0*p*_, *μ*_1*p*_, *μ*_2*ph*_, *μ*_3*pr*_) denotes the random coefficients for protein *p*; *μ*_4*h*_, *μ*_5*r*_, *μ*_6*k*_ denote the regression coefficients for the interactions terms of m/z ratio and experimental factors label and run respectively, they are not assumed to vary across proteins; *e_ilp_* denotes the unexplained residual error for peptide *i*, protein *p* and subject *l.*

At this level of the model, the intercept can vary across different proteins. The m/z ratio, runs and labels can have different effects on the peptide relative abundance across different proteins, and the intercept can vary across different biological samples.

### Define the protein and subject parts of the 2-level model

a. The regression coefficients ***U*** *=* (*μ***_0*p*_**, *μ***_l*p*_**, *μ***_2*ph*_**, *μ***_3*pr*_**) at the protein level can be described as the second level of the model,

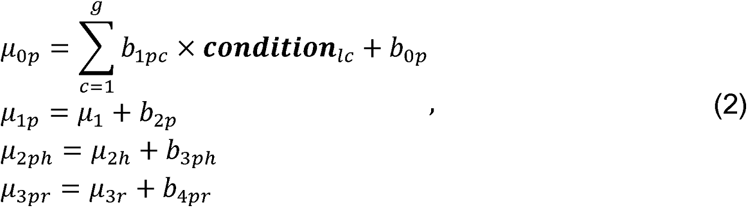

where ***condition****_lc_* represents a vector of binary variables to identify the physiological condition *c* ranged between 1 and *g* for subject *l*. *b*_0*p*_, *b*_2*p*_, *b*_3*ph*_, *b*_4*pr*_ represents the random residuals of protein *p* for the random intercept, m/z slope, label and run respectively; *b*_1*pc*_ represents a random residual of protein *p* and condition *c*. *μ*_1_, *μ*_2*h*_, *μ*_3*r*_ are the fixed intercepts. The protein part of the level 2 model defines the random regression coefficients of intercept, m/z ratio, label, run and subject’s physiological condition. It is assumed that they are different across proteins. There are no fixed terms for the physiological conditions at the protein level as they are separately defined at the subject level in equation (3).
b. The random intercepts at the subject level can be described separately as

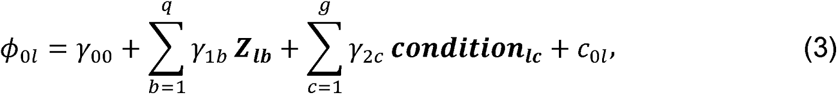

where *q* covariates of the subjects are represented by *Z =* [*Z*_*l*1_, *Z*_*l*2_…*Z_lq_*]; *γ*_00_, *γ*_1*b*_, *γ*_2*c*_ represents the fixed intercept and the fixed effect coefficients. *c*_0*l*_ represents the residual terms for subject *l*. The subject part of the level 2 model defines the random intercept for each subject. Let ***B*** be the vector of random effects for proteins, ***C*** be the vector of random residuals for subjects, and ***e*** be the vector of the random errors. We define the distribution of these random effect vectors as follows:

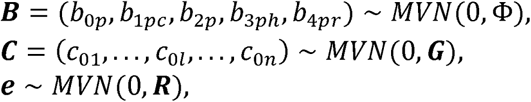

where Φ denotes the variance-covariance matrix for parameters at the protein level, ***G*** denotes the variance-covariance matrix for parameters at the subject level, and ***R*** is the variance-covariance matrix for the random residual errors at the peptide level. We assume the random effects for subjects are independent of the random effects for the proteins, and are independent of the random errors at the peptide level. The variance-covariance matrix of responses can be decomposed as ***V*** = K*_n_*_×*z*_Φ*_z_*_×*z*_K′*_z_*_×*n*_ + ***Z****_n×q_**G**_q_*_×*q*_***Z***′*_q×n_* + ***R****_n×n_*. K*_n_*_×*q*_ is the random effect matrix for protein and ***Z****_n×q_* is the random effect matrix for subjects, *n* is the total number of intensity observations in the data set.

## 3. A mixture model to handle missingness mechanism - a Bayesian approach

After introducing the multivariate model (Equations (1-3)) for complete data, the missingness mechanism is integrated with the same inference model within a Bayesian framework.

### 3.1. The configuration of the inference model and the imputation model for missingness

In a labelling mass spectrometry experiment, two types of missing peptide data from MS platform are observed; one is identified as zero and the other is identified as blank. The zeros are the left-censored values lower than the detectable limit, and the blanks are missing intensities due to weak signals.

Through the Bayesian framework, we can learn the missing data information from the other studies and incorporate them through the priors of the model. For example, the missing probability of the intensity can be estimated using its associations with the instrumental features of mass spectrometry reported in the other studies. We firstly define the Bayesian multivariate multilevel inference model (4), and secondly define the missing mechanism given by equation (5). Equation (4) describes a two-level hierarchical model that is a special scenario of equation (1) without interactions. The complete intensity data ***y*** = (*y*^obs^, *y*^miss^, *y^censored^*) is assumed to be multivariate normal distributed with mean ***y****_i,p_* and variance-covariance matrix Ω*. y_ilp_*^obs^, *y_ilp_*^miss^, *y_ilp_^censored^* represents the observed, missing and censored log-transformed intensity values, of protein *p* and the subject / respectively.

**The inference model:**

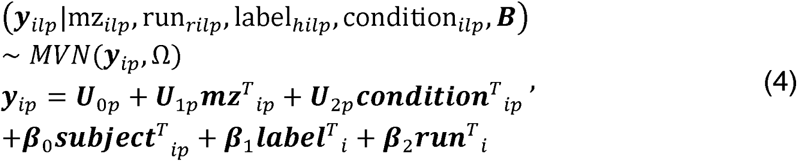

Where ***B*** *=* (***U***_0*p*_, ***U***_1*p*_, ***U***_2*p*_, ***β*_0_**, ***β*_1_**, ***β*_2_**), is the matrix of regression coefficients. ***U***_0*p*_, ***U***_1*p*_, ***U***_2*p*_ represent vectors of regression coefficients for different proteins. ***β*_0_**, ***β*_1_**, ***β*_2_** represent vectors of regression coefficients for different subjects, labels and runs respectively; they are assumed to be the same across different proteins. Ω is a variance-covariance matrix having a common unknown variance term *σ*^2^ across diagonal terms. ***y****_ip_* denotes the mean log-intensity for subject ***l***, ***y****_ip_* is predicted by the covariates of peptides, proteins and subjects in equation (4). 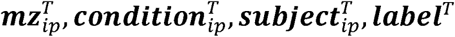 and ***run****^T^* represent the row vectors of explanatory variables m/z, physiological condition, subject, label and run respectively. Prior of the protein level regression coefficients (***U***_0*p*_, ***U***_1*p*_, ***U***_2*p*_) is multivariate normal distributed with mean ***α*** *=* (*α*_0*p*_, *α*_1*p*_, *α*_2*p*_) and variance-covariance matrix Σ, assuming Σ is the same for all subjects; The hyper-prior of ***α*** is multivariate normal distributed with mean ***f*** and variance-covariance ***G***.

A well-known hyper-prior for Σ is inverse-Wishart distribution[27], Σ^−1^ ~ Wishart(***R***, *d* = 3, *v*_0_), where R is a positive scaled definite matrix and *v*_0_ > *d* − 1. We can assign non-informative or informative hyper-priors for vector ***f***, matrices ***G*** and ***R***. Priors for the residual variance *σ*^2^ and the random intercept for subject ***β***_0_ and regression coefficients ***β*_1_**, ***β*_2_** are given as follows:

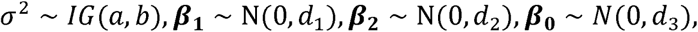

where *σ*^2^ is inverse-Gamma distributed with shape *a* and scale *b*, ***β***_0_, ***β*_1_**, ***β*_2_** are normal distributed, non-informative priors can be assigned to *a*,*b*,*d*_1_,…*d*_3_ if no prior information is available.

### 3.2 The missing mechanism

Let ***r*** be a binary indicator variable denoting if the intensity has a missing value (1: missing, 0: not-missing). Based on observations [Figure 1] and prior information, we know that the probability of missingness associated with the intensity value and the m/z ratio. If we assume that the probability of intensity being missed (including censored) is ***pm*** = {pm_*ilp*_} (for *r_ilp_* = 1), and it is Bernoulli distributed. The missing data mechanism is defined in the following equation (5):

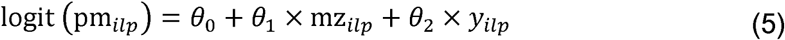

Let ***θ*** = (*θ*_0_, *θ*_1_, *θ*_2_) be the vector of missing data mechanism parameters. The conditional posterior distribution of *θ*_0_, *θ*_1_, *θ*_2_ is postulated as follows:

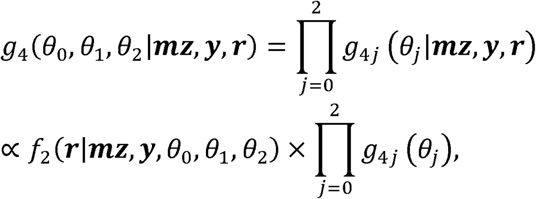

where ***mz*** *=* {mz_ilp_} represents the vector of m/z ratios, *g*_4*j*_ represents the density function for the priors of *θ_j_*(*j =* 0,1,2). *f_2_*(***r***|***mz***, ***y***) is the logistic regression likelihood function of equation (5). Because the missing peptide intensity is not observable in the data set, and no information can be obtained from the observations, a pair of informative priors is used based on the monotone relationship expected between missingness and intensity.

The censored values *y_ilp_^censored^* are proposed to belong to a truncated normal distribution with an upper threshold of *c*, *y_ilp_^censored^* ~ *N*(***y***^*censored*^|*c*).

### 3.3 Hamiltonian Monte Carlo (HMC) and Non-U-Turn posterior sampling (NUTs) algorithm

Our proposed method utilizes Hamiltonian MC/Non U Turn Sampling [28] for the posterior distribution. Compared to the Gibb sampling, Hamiltonian MC (HMC) [29] avoids the random walk by introducing the leapfrog function. It provides an alternative to approximating the solution on the continuous time scale from the solutions on the discrete time scale, with a specified step size. The logarithmic posterior probability function was simulated by one of the paired partial differentiated equations, namely the Hamiltonian. Larger moving steps are generated from the leap frog scheme, and this helps to improve the convergence compared to the random walk. It has been shown to have higher efficiency in sampling high-dimensional correlated multivariate distributions.

RStan is a new tool recently developed to implement HMC modelling for Bayesian data analysis, of which the posterior distribution can be sampled using the No-U-Turn Sampling method (NUTS)-an extension of HMC. NUTS [28] implements a recursive algorithm that will enable auto-tuning of the coupling parameters in HMC: numbers of leap-frog steps and the discrete step. In RStan, a known data variable cannot have missing values. If a variable contains missing values, the missing values will need to be defined separately as an unknown variable. The RStan algorithm and program for the model described in equation (4-5) are given in 3.4.

### 3.4 The program of the RStan HMC/NUTS for the missing data model (with only label as the covariate in this demonstration)

The RStan program for the proposed model comprises five blocks: data, transformed data, parameters, transformed parameters and model.

#### data

Step 1: Assign variables names to all the observed peptide intensities and their corresponding explanatory variables in the data block, excluding records with censored and missing intensities;

Step 2: Assign different variables names to all the observed explanatory variables linked to the censored and missing intensities in the data block;

Step 3: Assign priors and hyper priors of the covariance matrix for protein level covariates, and other known values (i.e. number of subjects, number of explanatory variables) in the data block.

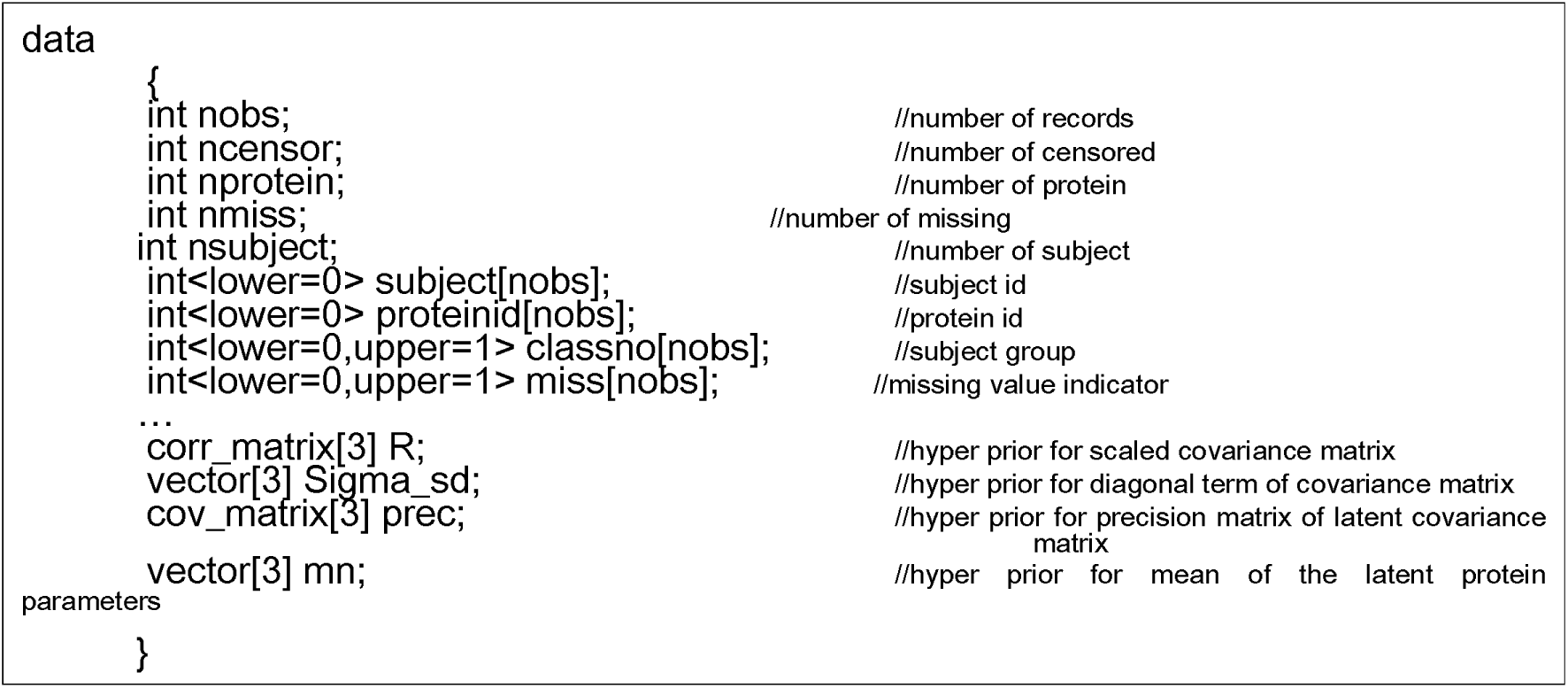

#### transformed data

Any data needing to be transformed in the program is given in the transformed data block. For instance, to derive a prior for the covariance matrix if the prior is given as a precision matrix in the data block.

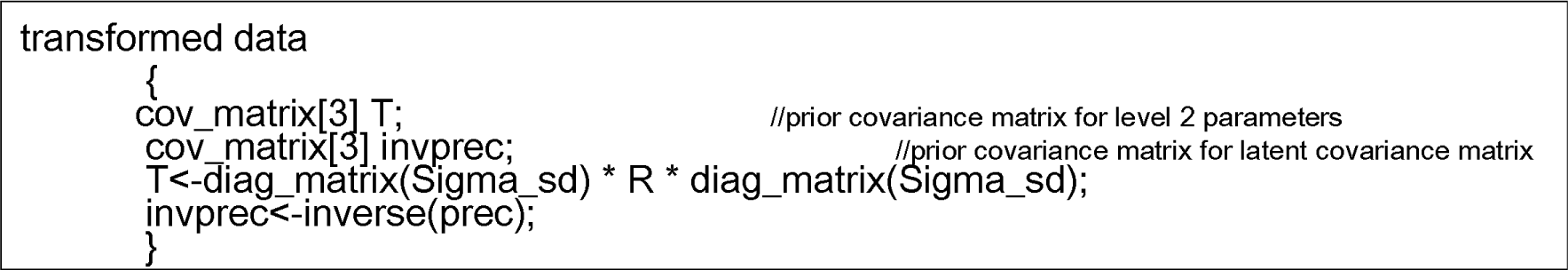

#### parameters

Any unknown variables including the missing and censored intensities, and variables that will be used to **re-parameterize** the unknown variables, are defined in the parameter block. The re-parameterization is to improve the statistical efficiency in posterior samplings[30].

Variables used for re-parameterizations are named as latent variables. In the NUTS program in the supplementary material, the latent parameters defined in this block are named by adding “latent” to the unknown parameters. For example, beta4_ latent is the latent variable for regression coefficient beta4.

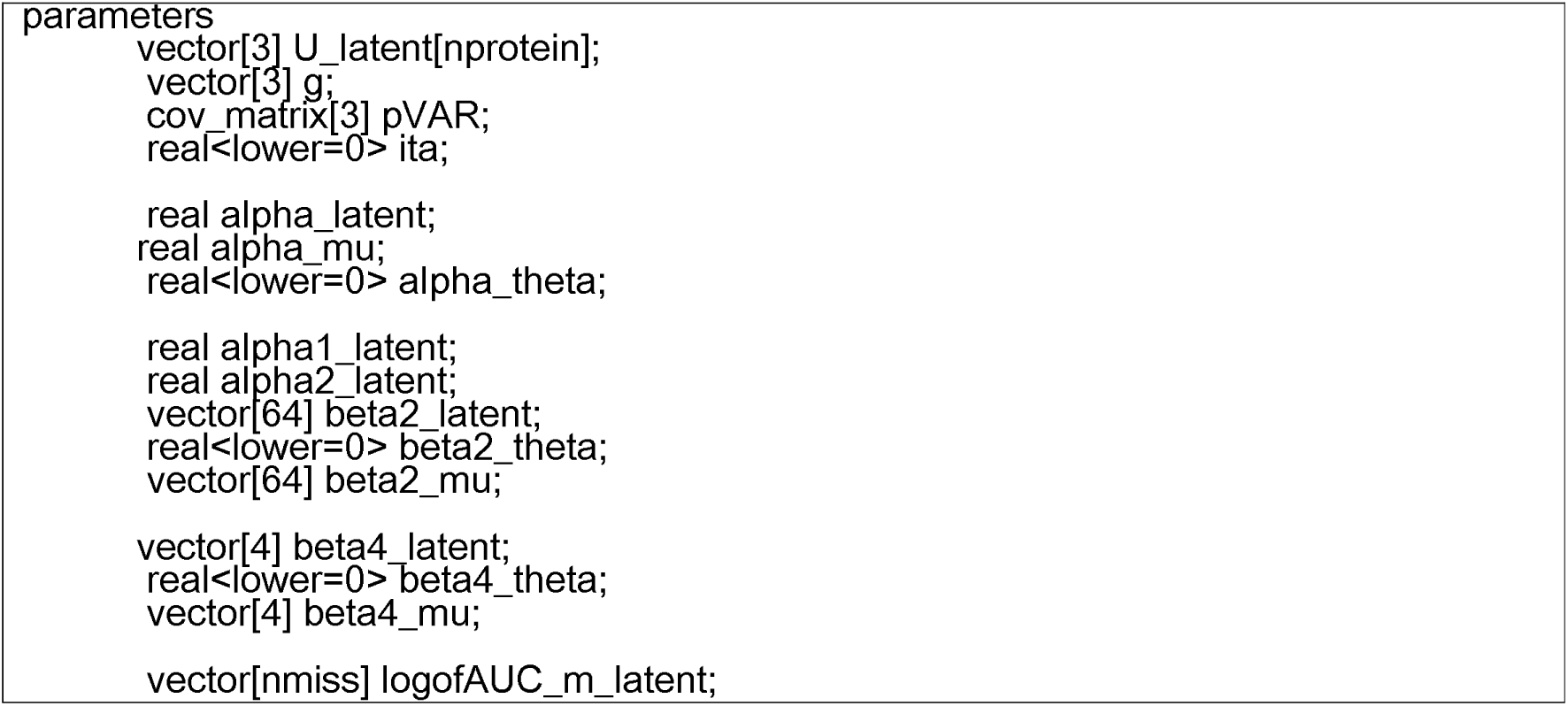

**<u>transformed parameters</u>**

Any unknown variable that comprises both known and unknown values can be derived in the transformed parameters block. The missing and censored peptide quantities and the probability of missing are derived in this block. The parameters use re-parameterization techniques are also derived from this block.

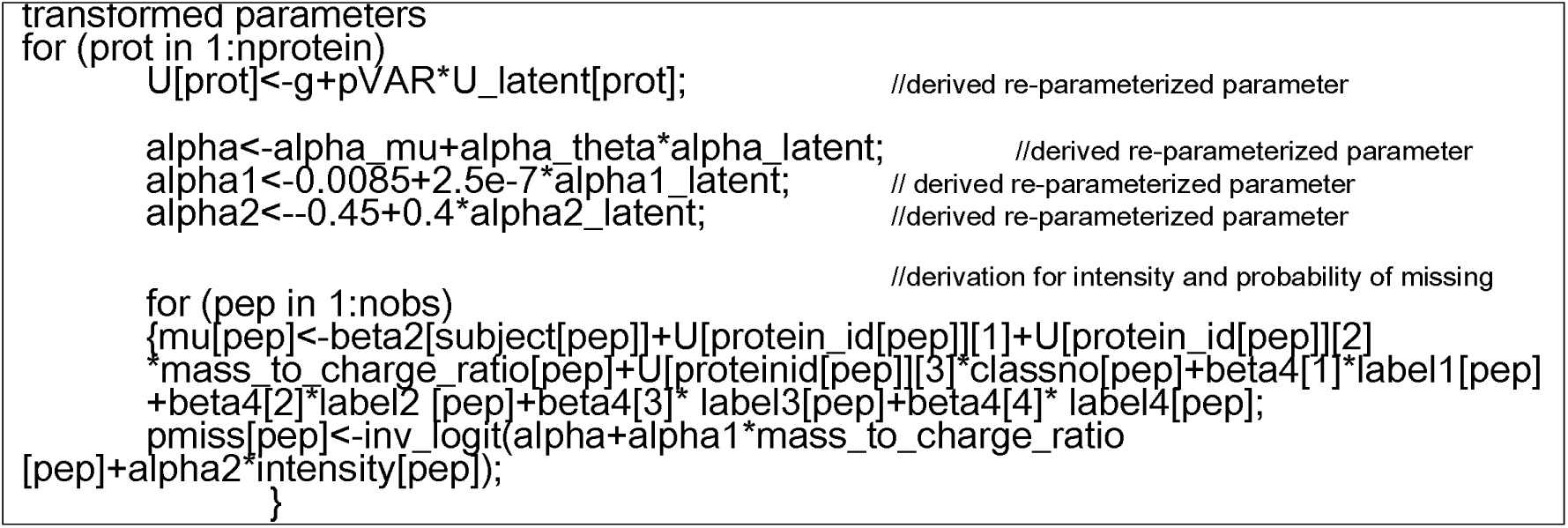

#### model

In the model block, all the unknown parameters are defined by its sampling distribution using ~.

Step 1: Sampling for the define priors and hyper priors: distributions of all priors or hyper priors for unknown parameters are defined firstly in the model block.

Step 2: Sampling for the latent variables used for re-parameterization: the re-parameterization is equivalent to a two-steps sampling that firstly samples the latent parameters (i.e., location and scale) according to their hypothesized distributions in the model block; and secondly derives the unknown parameters using the selected latent parameters in the transformed parameters block.

Step 3: Missing and censored data parameters: The missing and censored values are treated as a couple of unknown variables. Logistic regression is used to model the missing mechanism and is defined in the transformed parameter block. The posterior distribution of missing values is sampled through the re-parameterization. The posterior distribution of the censored value is estimated by integration of a truncated normal distribution.

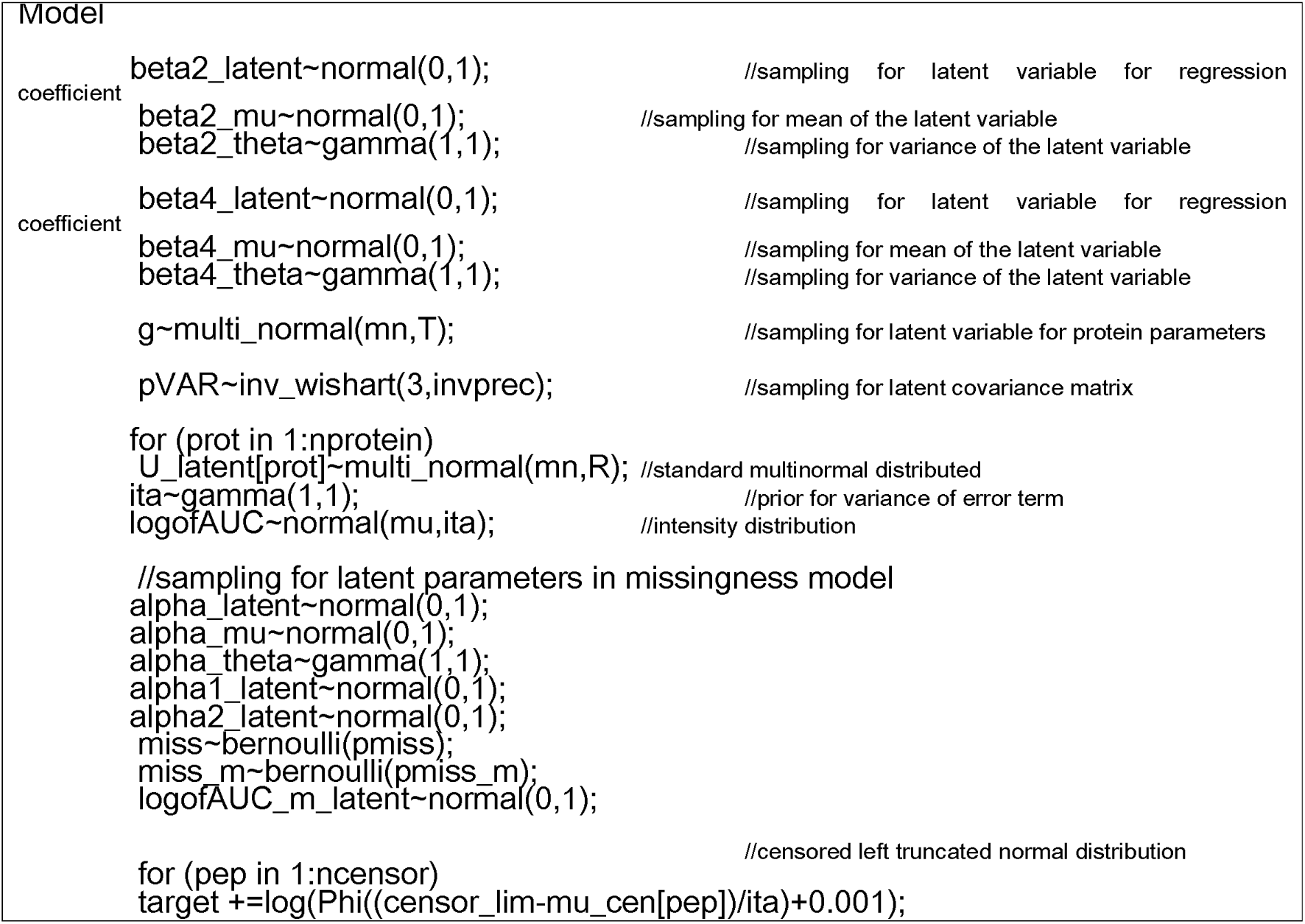

## 4. Using simulation to validate the model for the missing mechanism and compare between *lmer*, BUG and NUTs estimation

An iTRAQ proteomic experiment with complete data was simulated; the same dataset with non random missing patterns as described in [21] was added in the same dataset. The estimates from three different methods (lmer, BUG and NUTs) were compared against the known parameters from the complete data.

The simulated study has a row-column design for eight runs (row), eight labels (column) and two classes that comprised 32 subjects from the healthy population as controls and 32 subjects from the diseased population. A Poisson distribution (*λ*=5) was used to generate the number of peptides of 200 proteins. The intensity of the peptide has confounding effects of run, label, the total amount of protein in fluid tissue sample, subject class, protein intercept and mass-to-charge ratio (m/z). The non-random missingness pattern are associated with the peptide intensity and the mass-to-charge ratio (m/z). The censored threshold was set to 0.1 which was the minimal log-intensity of an iTRAQ case study. Any simulated log-intensity below this value was set to zero. The simulated data set of 78400 records has 928 (1.2%) observations with censored peptide intensities and 14287 (18.2%) observations with completely missing peptide intensities.

We compared the univariate ANOVA, a multivariate mixed model fitted with *lmer* in R [31] using complete data only; we also compared the multivariate *lmer* estimates with the MNAR Bayesian model estimates that including observed covariates of missing intensities.

Using complete data, estimates of mean class differences (diseased vs. control) from the multiple proteins R/*lmer* model demonstrated a better agreement with the actual values, than estimates from the univariate single protein models did (scatter plots figure 2). In two sets of similar linear regressions that include actual values as dependents, estimates of single or multiple protein models are included as independents respectively comparing against the actual values. The R^2^ is 0.87 with a slope of 1.21 (standard error: 0.034) in the regression using estimates from the univariate single protein model. The R^2^ is 0.91 with a slope of 1.27 (standard error: 0.028) in the regression using estimates from the multiple proteins mode. The R^2^ is larger and standard error of the slope is slightly smaller using estimates from the multiple proteins model.

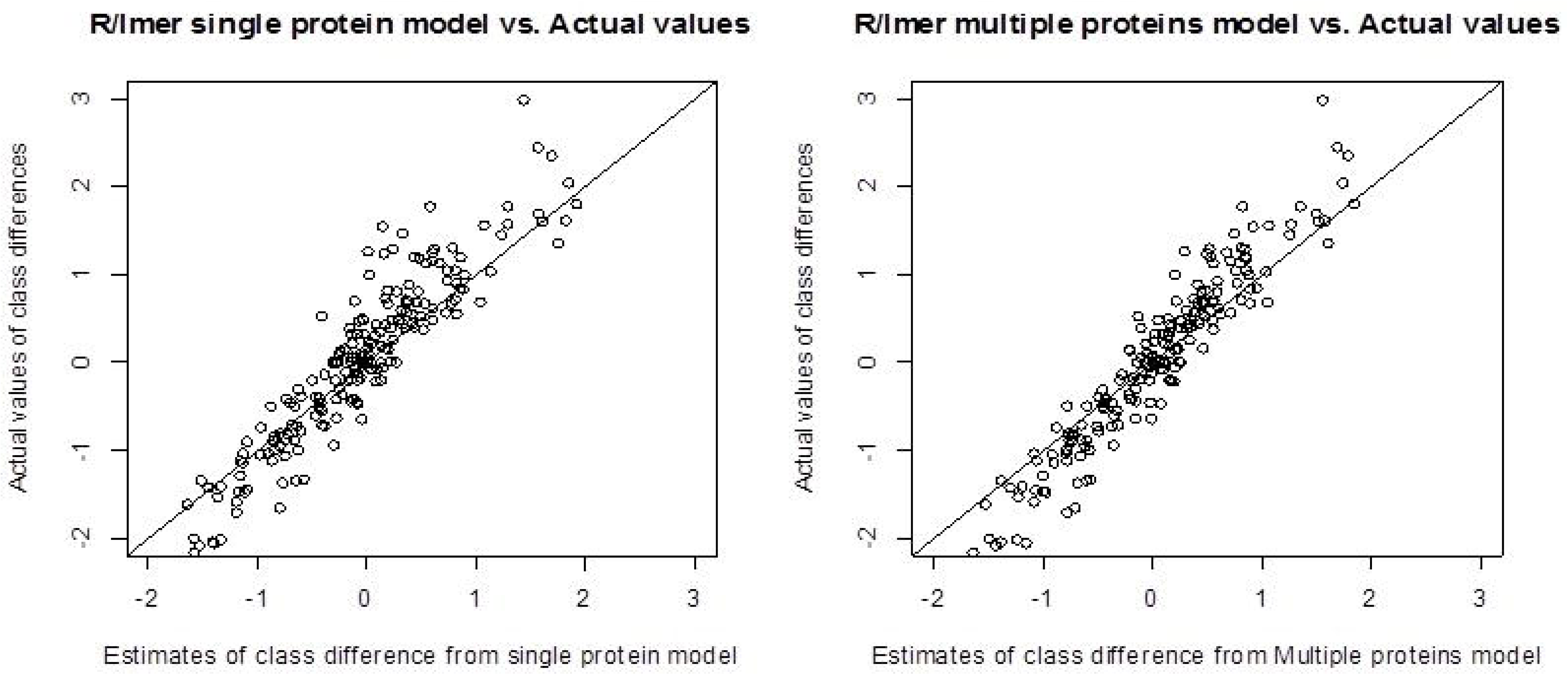

When the non-random missingness mechanism is modelled using the Bayesian method, a program using BUGS and a program using HMC/NUTS were used to derive the posterior distributions of unknown parameters. Both programs have smaller variability in the estimates of the mean class difference in comparison to the actual values (scatter plot figure 3). The HMC/NUTS program, which models the censored value as normal truncated distributed, achieves a better agreement with the actual values. This is indicated by the closer-to-1 slope of the linear regression that has the HMC/NUTS estimates compared against the actual values. The achieved R^2^ Is 0.95, and the slope is 1.07 (standard error: 0.018).

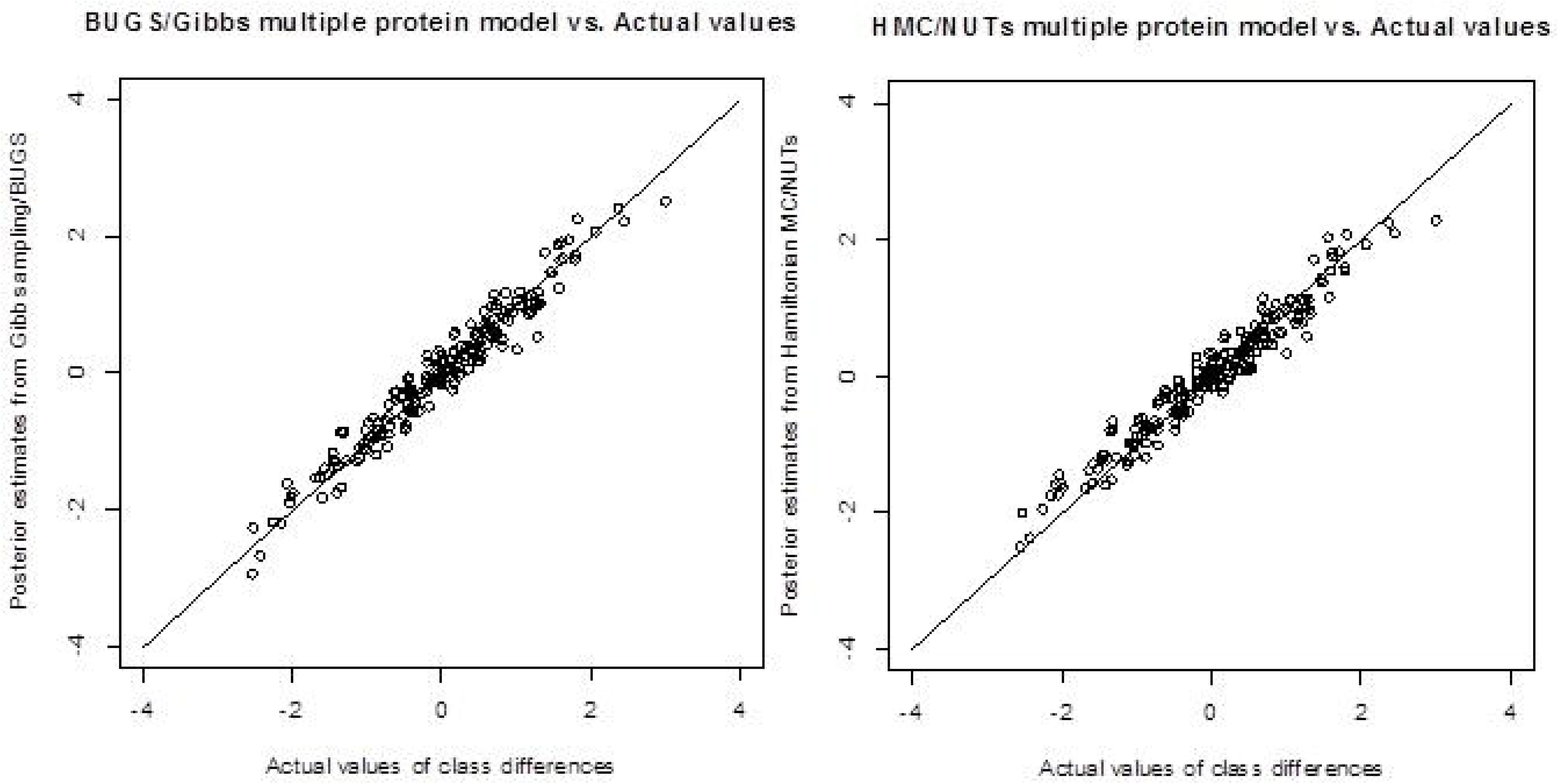

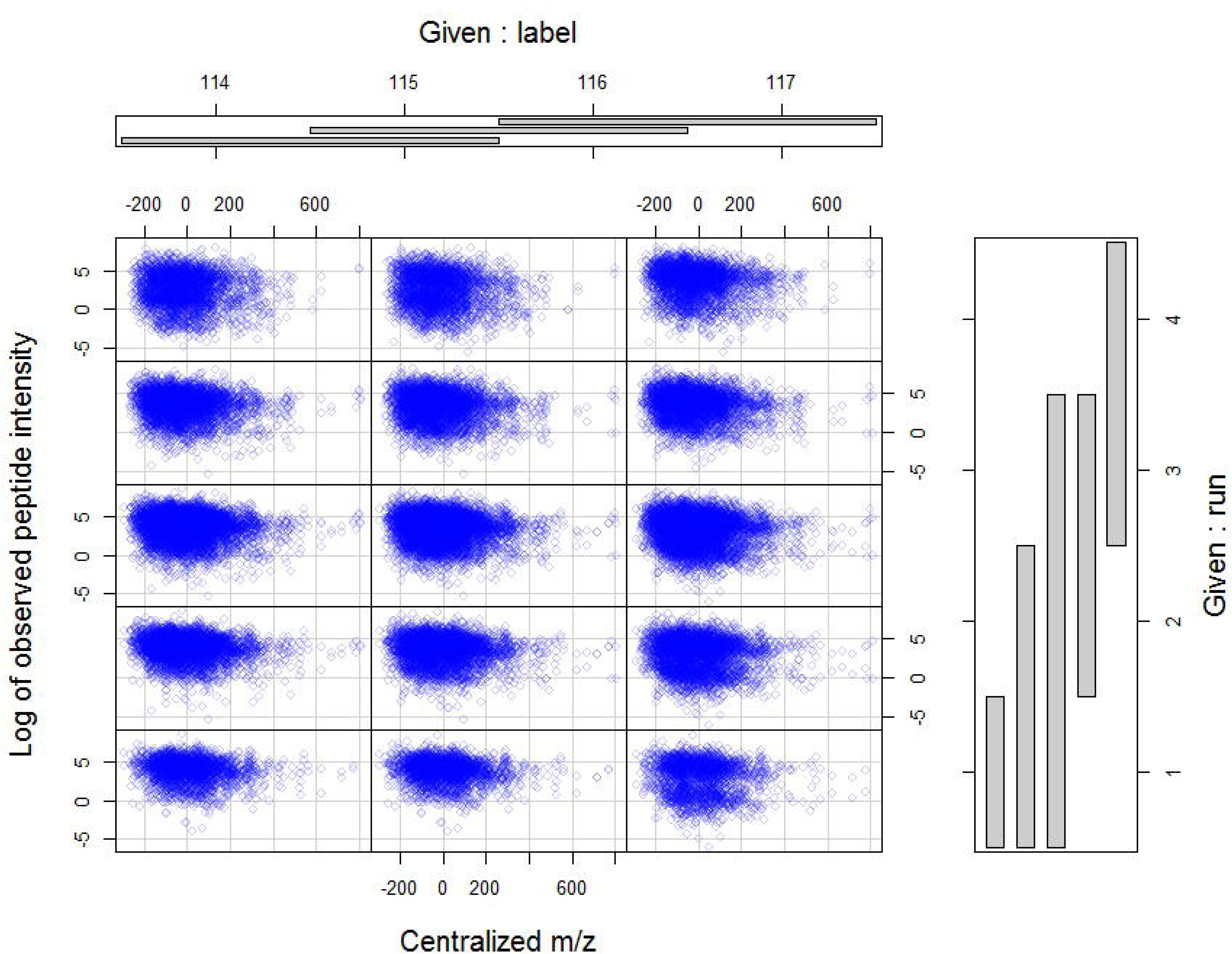

The unexplained residual variance of the model using the HMC/NUTS algorithm is 1.87 (95% credible interval: 1.81, 1.93), which is similar to the actual value of 2.0, and smaller than 3.11- the result of the R *lmer* model of the complete cases [Figure 3].

## 5. A cardiac proteomic case study

The cardiac proteomic study is one part of a double blinded randomized control trial [32] that investigated how the intra-coronary metoprolol (beta blocker) changed the myocardial metabolism of peptides, proteins and metabolites profiles in patients admitted to hospital with a first myocardial infarction. The cardiac proteomic study investigated how the whole plasma proteome profile changed after coronary angioplasty intervention. The study collected five plasma samples at different time points in eight patients during their percutaneous coronary interventions. Two of these plasma samples (before and 20 minutes after a controlled coronary occlusion) were selected in the proteomic study. The IPI human database (European Bioinformatics Institute, Hinxton, UK) was used in the Protein Pilot Software to match the observed peptides with their corresponding proteins for the identification.

This case study has 17780 peptide observations of which 1667 (9.4%) had censored and 218 (1.2%) had completely missing values in the peptide intensities. The Bayesian model with the selection model for the missing mechanism is constructed as follows:

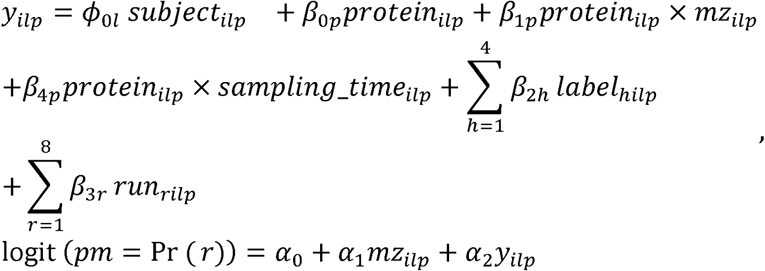

where *y_ilp_* defines the mean for the peptide intensities including complete, completely missing and censored values on the log scale, *y_ilp_* ~ *N*(*μ_p_*, *σ*^2^). *subject_ilp_* defines the subject identity and *ϕ*_0*l*_ defines the intercepts for each subject. *β*_0*p*_, *β*_1*p*_, *β*_4*p*_ define the protein level parameters: intercept, m/z ratio and sampling time. The difference in log-intensities between sampling time is equivalent to the intervention effect. *protein_ilp_* is a variable recording the identity of protein *p*. *β*_2*h*_ and *β*_3*r*_ are regression coefficients for label *h* and run *r* respectively. *pm* defines the probability of having a missing intensity value; and *α*_0_, …, *α*_1_ define the regression coefficients in the logistic regression for *pm*.

The coefficients of logistic regression *α*_1_ and *α*_2_ are part of the joint unknown parameters with a pair of chosen normal distributed informative priors *N*(0.0085,4 × 10^−8^), *N*(−0.45, 0.25) for m/z ratio and peptide abundance respectively [21]. The protein level parameters *β*_0*p*_, *β*_l*p*_, *β*_4*p*_ used a multivariate normal distributed prior (**y**, 7) where,

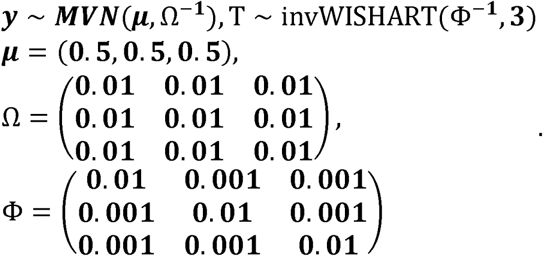

The non-informative priors for the peptide level parameters *β*_2,*h*_ and *β*_3,*h*_ are normal distributed and denoted as *N*(0,10) with mean 0 and precision 0.1, and the non-informative prior for the subject level parameters *ϕ*_0,*l*_ is normal distributed and denoted as *N*(0,1) with mean 0 and precision 1.

Using the complete data, the significant fixed effect of the intervention derived from the R/*lmer* model reveals a systematic shift of -1.42 in all protein expressions on the log-scale; the other significant fixed effects include the centralized m/z, label 116 and 117 (table1.). The predicted random effects of intervention for different proteins indicate the magnitudes of fold changes in the protein abundances introduced by the intervention (figure 5). Compared to the variance components in R/lmer model, using NUTS model results in a similar intervention variance but a larger m/z ratio variance. The unexplained residuals variance in the NUTS and BUGS models are 1.62 and 0.37 respectively (table 2); both are smaller than 2.57 in the R/*lmer* multiple protein models.

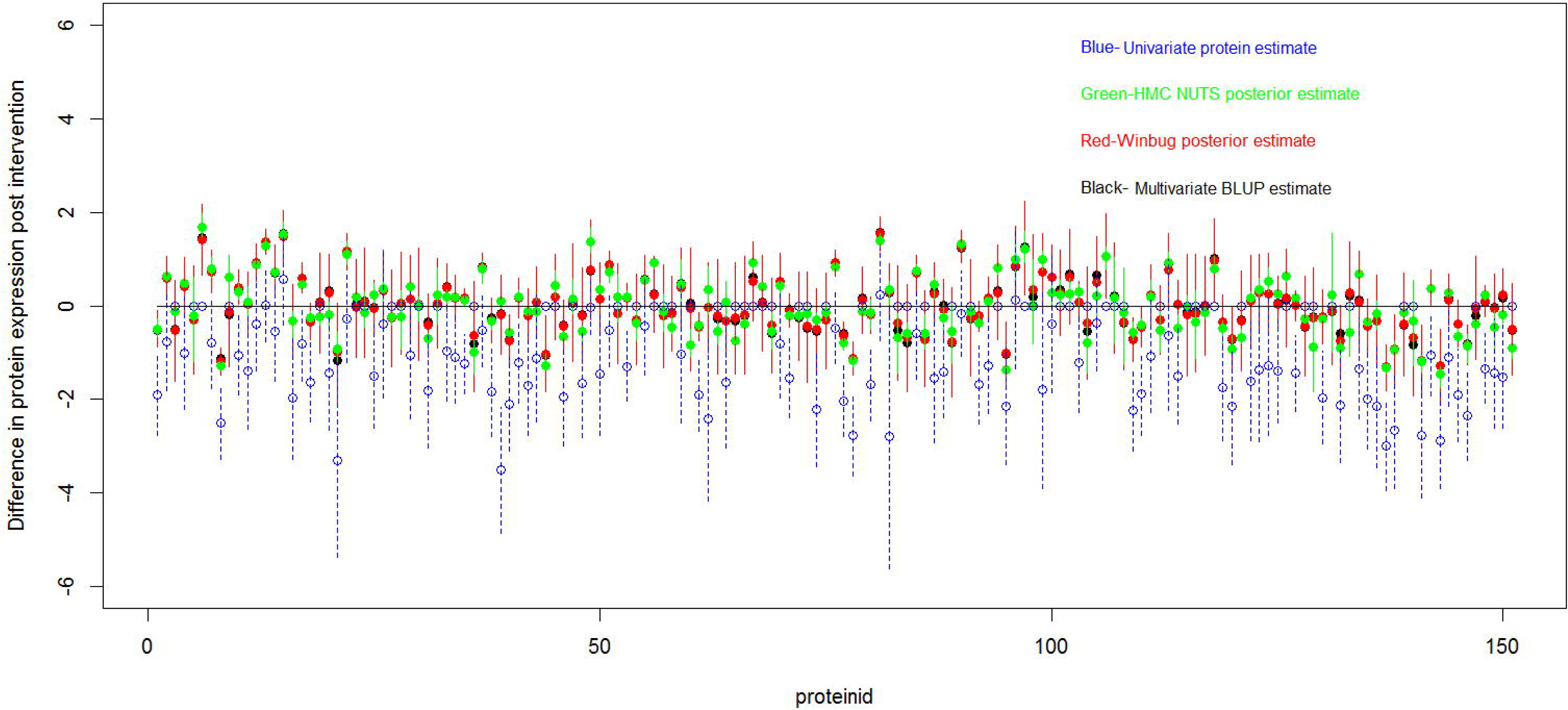

## 6. Discussion

The multilevel model proposed in this paper provides a new approach to analysing a group or all groups of proteins when protein abundances comprise missing values that are Missing-Not-At-Random. The flexibility of multivariate multilevel model allows for an unbalanced design structure among the responses. This becomes an advantage in a proteomic study when the numbers of component peptides of the same protein are unequal across different subjects. Within this multivariate framework, experimental factors of multiple proteins can be utilized in conjunction with clinical design factors to derive the results of a single protein. Inclusion of covariates from multiple proteins improves precision of the parameters at protein level and results in shrunk estimates for a single protein with few observations. The multivariate model also enables identification of proteome-wide characteristics, such as systematic differences in protein abundances between two sampling times in the cardiac case study.

When non-random missingness needs to be considered and modelled, using numerical integration or an EM algorithm is not feasible in such a complex framework. Using Hamiltonian MC/NUTS is an improvement, especially in the case of utilizing non-standard distribution for handling missing data. Compared to BUGS, RStan is more flexible for handling missing data via the truncated distribution function. The NUTS algorithm is more efficient in dealing with the truncated distribution and non-standard distribution than BUGS.

The philosophy of the proposed multilevel multivariate method can be generalized to other proteomics or “omic” research with different experimental structures. Future development of the model will include consideration of various covariance matrices for protein and subject level variables, and adding known factors as explanatory variables to estimate components of the covariance matrices.

## Acknowledgements

The authors express their gratitude to Dr Beatrix Jones who have given valuable comments to the study. The authors also acknowledge the investigators Dr Patrick Gladding and Prof Ralph Stewart for their contributions to the cardiac proteomic study; Greenlane Research and Education Fund for the cardiac proteomic study; and New Zealand eScience infrastructure (NeSI) facilitated the program testing.

